# Biological Continued Pretraining Reshapes the Capability Profile of a Foundation Model Without Catastrophic Forgetting

**DOI:** 10.64898/2026.07.06.736700

**Authors:** Liang Wang

## Abstract

It is widely assumed that continued pretraining (CPT) on a narrow, out-of-distribution corpus such as raw biological sequence must trade away a general-purpose model’s broad competence — the “alignment tax” or catastrophic-forgetting intuition. We test this directly, without any new training, by re-analyzing three checkpoints from a single lineage of a 26B-parameter Mixture-of-Experts model (Gemma-4-26B-A4B): the instruction-tuned base, the same model after biological CPT (8.7B tokens of DNA, protein, and biomedical text), and after subsequent supervised fine-tuning (SFT). Across three independent capability axes — general knowledge/reasoning (MMLU, ARC, HellaSwag), code generation (MBPP), and biomedical knowledge (BixBench) — we find that biological CPT does *not* degrade the model; it **lifts** it: MMLU +13 points, MBPP pass@1 nearly doubles (0.33 →0.63), and BixBench discrimination rises sharply (MCC 0.23 → 0.92). The single measured regression is truthfulness (TruthfulQA 8.8 points), a small and interpretable domain drift. A clean vocabulary-expansion ablation (*<* 0.4 pt on every general metric) confirms the gains are attributable to CPT, not tokenizer changes. Crucially, subsequent SFT *narrows* the model back: all three axes fall to near-base levels, revealing a consistent division of labor — **CPT re-organizes and lifts the shared capability substrate; SFT cashes it out onto target tasks**. We argue this reframes biological sequence not as a competitor for a foundation model’s capacity but as a form of structured scientific data that reshapes its capability profile, and that CPT and SFT should be budgeted as complementary rather than substitutable stages. All checkpoints, evaluation code, and per-example outputs are public.

**Highlights:**

- A training-free re-analysis of one 26B MoE lineage isolates the effect of biological continued pretraining (CPT) from tokenizer changes and from fine-tuning.
- Biological CPT does not cause catastrophic forgetting; it *raises* general knowledge (MMLU +13 pts) and code generation (MBPP pass@1 0.33 *→* 0.63).
- CPT also makes chain-of-thought reasoning 41% shorter and near-backtrack-free while pre-serving accuracy — an effect invisible to accuracy metrics.
- A consistent CPT-lifts / SFT-narrows division of labor recurs across four axes, reframing biological sequence as structured scientific data that reshapes a model’s capability profile.

**The Bigger Picture:** Adapting a general-purpose AI model to a specialized domain — here, the language of DNA and proteins — is usually assumed to come at a cost: teach it biology and it forgets how to reason about everything else. This “no free lunch” intuition shapes how practitioners budget compute and whether they attempt domain adaptation at all. We test the assumption directly, and without running any new training, by comparing three snapshots of the same model taken before and after biological training. The result overturns the intuition: feeding the model raw biological sequence made it *better* at general knowledge, at writing code, and even changed *how* it reasons — producing shorter, more decisive chains of thought without losing accuracy. The gains appear during the sequence-pretraining stage and are partly given back during task-specific fine-tuning, revealing that the two stages play complementary rather than interchangeable roles. This suggests a broader principle for data-centric AI: structured scientific data — biological sequence today, and by extension code, mathematics, and chemistry — is not merely knowledge to be absorbed but a lever that reshapes what a foundation model can do.

## 1 Introduction

The dominant intuition about continued pretraining (CPT) on a narrow domain corpus is subtractive: pouring out-of-distribution tokens — raw protein and DNA sequence, which share almost no surface statistics with natural language — into a general-purpose model is expected to erode its broad competence, a phenomenon variously described as the *alignment tax* or *catastrophic forgetting* [1, 2]. Under this view, domain adaptation is a trade: one buys biological ability at the cost of general ability, and the practitioner’s job is to minimize the damage.

This paper argues that, for a modern Mixture-of-Experts foundation model, the intuition is wrong — or at least incomplete. We show, by *re-analysis of existing checkpoints and without any new training*, that biological CPT on a 26B-parameter model does not merely avoid catastrophic forgetting; it *improves* the model on capability axes that have nothing to do with biology, including general knowledge (MMLU [3]) and code generation (MBPP). We then show that the subsequent supervised fine-tuning (SFT) stage, far from adding capability, *narrows* the model back toward its base competence on held-out axes. The picture that emerges is a clean division of labor between the two training stages, and it suggests a reframing of what biological data *is* to a foundation model. Our substrate is a Mixture-of-Experts model [4–6] (Gemma-4-26B-A4B [7]); the analysis is deliberately training-free, re-using an existing CPT/SFT lineage [2] so that every capability difference is attributable to a single stage. This complements router-level interpretability of the same model family [8–10] by asking not *where* computation is organized but *how* the training data reshapes what the model can do.

### Related work

Our claim sits at the intersection of three literatures. *Continued pretraining and forgetting*: domain-adaptive CPT is standard practice [2, 11, 12], but is widely reported to risk catastrophic forgetting of general ability [1, 13], motivating replay and balancing schemes.

Most of this evidence is on dense models and measures forgetting as a cost to be minimized; we instead find a net *gain* on a sparse MoE model and attribute it to the structural regularity of the pretraining corpus rather than its factual content. *Biological foundation models*: protein and genomic models — ESM-2 [14], ProtT5 [15], DNABERT-2 [16], Nucleotide Transformer [17], the Evo family [18, 19], and cross-modal LucaOne [20], as well as LLMs adapted to biology [21–23] — are typically evaluated only on in-domain tasks, leaving open whether biological training helps or harms general competence. *Data-centric AI and scaling*: work on pretraining-data composition and its effect on capability [24, 25] argues that *what* a model is trained on shapes *what* it can do; we provide a controlled, single-lineage instance of this principle where the manipulated variable is a scientific-sequence corpus and the readout spans knowledge, code, and reasoning behavior. To our knowledge no prior work reports that biological CPT *improves* general and coding benchmarks, nor that it measurably alters reasoning style.

### Contributions

(i) A training-free re-analysis protocol over a single checkpoint lineage that isolates the effect of biological CPT from tokenizer changes and from SFT. (ii) Evidence that biological CPT lifts, rather than degrades, general and code capability, with a single interpretable regression (truthfulness). (iii) A consistent CPT-lifts / SFT-narrows division of labor observed across four independent evaluation axes. (iv) A data-centric reframing: biological sequence as structured scientific data that reshapes a foundation model’s capability profile.

### 2 Setup: one lineage, three checkpoints

Our analysis rests on a single training lineage, so that every capability difference we report can be attributed to a specific training stage rather than to confounds of architecture, tokenizer, or data version.

### Base model

All checkpoints derive from Gemma-4-26B-A4B, a Mixture-of-Experts transformer (128 experts per layer, top-8 routing, ≈ 3.8B active parameters). We use the instruction-tuned release as the origin of the lineage; the biological CPT and SFT checkpoints are those of OmniGene-4 [8].

**The three (four) checkpoints**. The lineage is

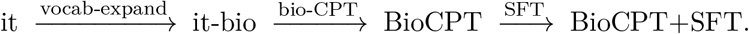

**(1) Base (it):** the original instruction-tuned model, vocabulary 262,144. **(2) Base (it-bio):** identical weights with the vocabulary expanded to 290,172 by adding 28,028 biological BPE tokens (DNA/protein), but *no further training*; this serves as a clean control that isolates the cost of tokenizer expansion alone. **(3) BioCPT:** it-bio after continued pretraining on 8.7B tokens of mixed biological corpus (DNA, protein sequence, and biomedical literature text), with LoRA (*r*=64) over attention, MLP, and the MoE router, embeddings unfrozen; merged to full weights. **(4) BioCPT+SFT:** BioCPT after supervised fine-tuning on biological instruction data (a dual-head structural variant, “v5”); merged to full weights. We verified the lineage by direct weight comparison: it and it-bio share router/expert weights bit-for-bit (mean abs. diff. 0), differing only in embeddings, while BioCPT’s router and expert weights have measurably moved.

#### Evaluation

We use the `lm-evaluation-harness` (v0.4.12) for all general and coding benchmarks, at community-standard few-shot settings (MMLU 5-shot, ARC-Challenge 25-shot, HellaSwag 10-shot, TruthfulQA-mc2 0-shot; HumanEval 0-shot, MBPP 3-shot). Multiple-choice tasks are scored by loglikelihood, which is robust to a model’s chat-vs-completion output format — a property we exploit deliberately, since our lineage mixes an instruction-tuned base with a completion-style CPT checkpoint. Biomedical discrimination (BixBench) is likewise scored by loglikelihood of the answer choices, and reasoning-behavior diagnostics (§3) use a shared 8-shot chain-of-thought scaffold. All runs are on a single 96 GB GPU in bfloat16; the full model (52≈ GB) fits on one card, and the entire study required no training.

#### Honest scope

Two evaluation regimes are format-sensitive and we flag them up front. HumanEval 0-shot (bare function-completion) is unfair to instruction/SFT checkpoints and we exclude it from conclusions, reporting MBPP 3-shot as the fair coding metric. Zero-shot protein *homology* is not measurable by any simple probe we tried (forced Yes/No loglikelihood is near-random for all checkpoints; free generation is format-confounded); we therefore treat sequence-homology ability separately and do not force it into the transfer analysis.

## 3 Results

The central finding is a single pattern that recurs across every capability axis we measured: biological CPT *lifts* the model, and subsequent SFT *narrows* it back (Figure 1, Figure 3). We present the three completed axes, then the reasoning-behavior diagnostics.

**Figure 1.**
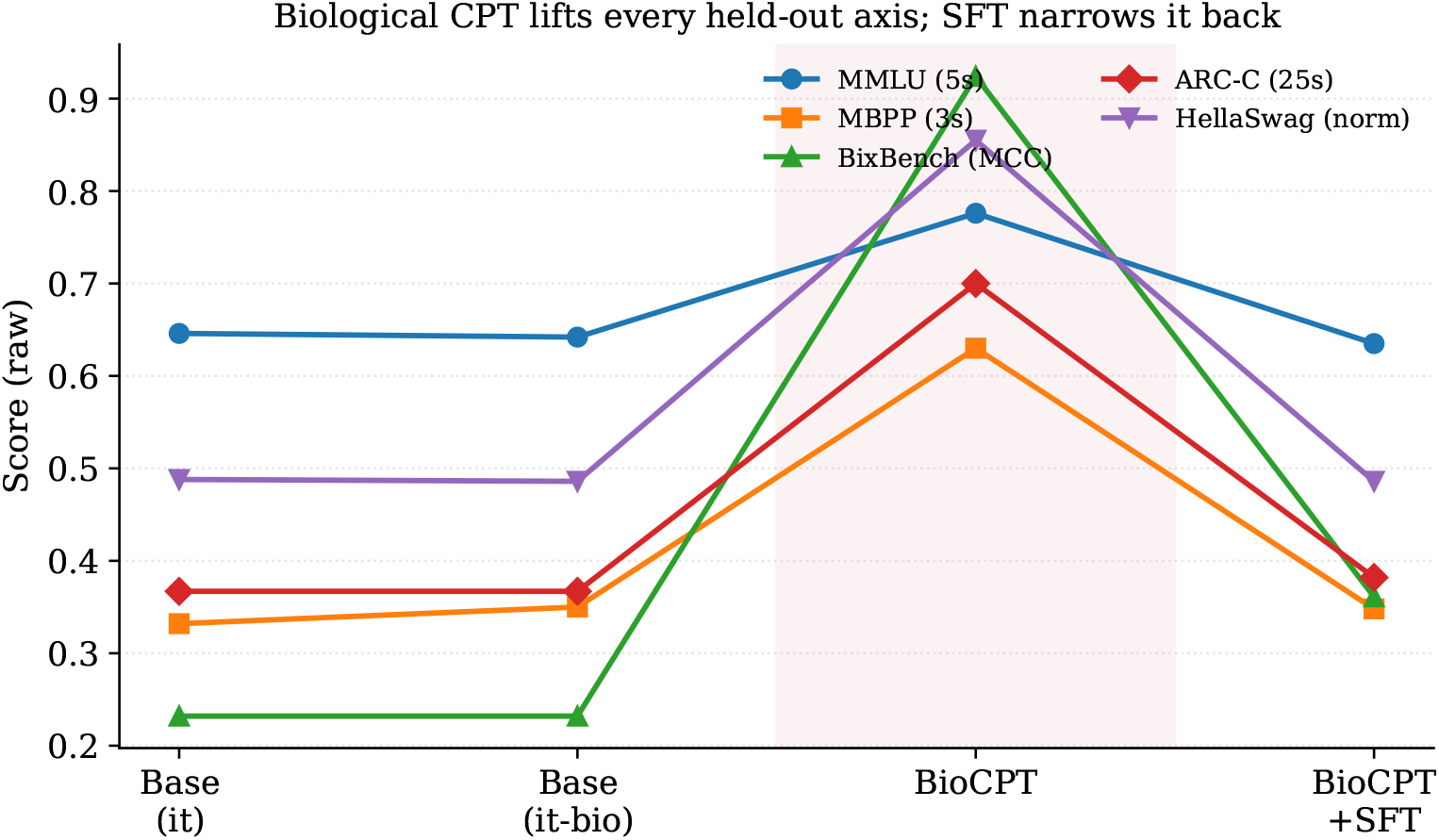
Every held-out capability axis follows the same trajectory: flat across the vocabulary-expansion control (it → it-bio), a sharp rise at biological CPT (shaded), and a fall back toward base after SFT. Raw scores; see Tables 1–2.

### 3.1 Vocabulary expansion is free

Before attributing anything to CPT we rule out the tokenizer. The original model (it) and the vocabulary-expanded but untrained model (it-bio) are indistinguishable on every general benchmark (MMLU 0.646 vs. 0.642; ARC-C 0.367 vs. 0.367; HellaSwag-norm 0.488 vs. 0.486; TruthfulQA 0.533 vs. 0.532; all ≤ 0.4 pt). Adding 28K biological tokens to the vocabulary therefore costs nothing in general competence, and any change at the BioCPT stage is attributable to the continued pretraining itself.

**Table 1.** (1) General. Biological CPT lifts general knowledge/reasoning; SFT narrows it back. Loglikelihood multiple-choice; community-standard few-shot.

| Checkpoint | MMLU (5s) | ARC-C acc (25s) | ARC-C norm | HellaSwag norm (10s) | TruthfulQA-mc2 |
| --- | --- | --- | --- | --- | --- |
| Base (it) | 0.646 | 0.367 | 0.403 | 0.488 | 0.533 |
| Base (it-bio) | 0.642 | 0.367 | 0.402 | 0.486 | 0.532 |
| **BioCPT** | **0.776** | **0.700** | **0.730** | **0.855** | 0.445 |
| BioCPT+SFT | 0.635 | 0.382 | 0.396 | 0.486 | 0.562 |

### 3.2 General knowledge and reasoning rise, not fall

Table 1 shows the (1) General axis. Against the intuition of catastrophic forgetting, biological CPT *improves* the model: MMLU +13.0 points (0.646 → 0.776), ARC-Challenge accuracy nearly doubles (0.367→0.700), and HellaSwag-norm rises +36.7 points. The one regression is TruthfulQA (−8.8 points), an interpretable drift on truthfulness/anti-misinformation after heavy exposure to sequence and literature text. We note honestly that part of the ARC/HellaSwag jump is mechanistic — an instruction-tuned model is weak at raw-loglikelihood multiple choice, and CPT on raw text restores base-model-like MC behavior — but MMLU (+13 pt, knowledge-heavy) is a hard gain unaffected by this. Subsequent SFT returns all four metrics to near-base levels.

### 3.3 Code capability nearly doubles

Code is a structured formal language, and the (3) Coding axis (Table 2) shows the same shape. On MBPP (3-shot, the format-fair metric), pass@1 rises from 0.33 at base to **0.63** after BioCPT, then falls to 0.35 after SFT. (HumanEval 0-shot is excluded: bare completion is unfair to instruction/SFT checkpoints, which score ≈0; we report it only for completeness.) That biological-sequence CPT nearly doubles Python coding ability is consistent with the hypothesis that structured-sequence pretraining strengthens a shared substrate for formal/structural languages.

**Table 2.** (3) Coding. MBPP 3-shot is the fair metric (HumanEval 0-shot excluded, see text).

| Checkpoint | HumanEval 0s (excl.) | **MBPP 3s pass@1** |
| --- | --- | --- |
| Base (it) | 0.012 | 0.332 |
| Base (it-bio) | 0.000 | 0.350 |
| **BioCPT** | 0.439 | **0.630** |
| BioCPT+SFT | 0.000 | 0.348 |

### 3.4 Biomedical knowledge: the same curve

On BixBench (biomedical true/false, loglikelihood-scored MCC), the (2) Biology axis reproduces the pattern exactly: MCC 0.232 at base →**0.924** after BioCPT →0.361 after SFT. Protein sequence-homology, by contrast, is not measurable zero-shot by any simple probe (forced Yes/No loglikelihood is near-random for all checkpoints; see §2); we read this as evidence that sequence-homology ability is *installed into representations* by CPT but only *cashed out* into task performance by SFT, consistent with prior dual-mode results on this model family [8].

### 3.5 Reasoning behavior: CPT makes the model concise and decisive

The three axes above concern *what* the model knows. We finally ask whether CPT changes *how* it reasons, independent of accuracy. On 200 GSM8K problems, each sampled *k*=5 times under a shared 8-shot chain-of-thought scaffold, we measure chain length, self-correction markers (“wait/actually/however/reconsider”), hedging markers, and self-consistency (agreement of the final answer across the 5 samples). Table 3 and Figure 2 show a striking behavioral shift at the BioCPT stage: the reasoning chain *shortens by 41%* (108.8→64.4 tokens) and self-correction *nearly vanishes* (0.370→0.006 backtracks per generation), yet accuracy does not fall (it rises slightly, 0.667→0.689) and self-consistency is unchanged (0.745). Biological CPT — pretraining on deterministic, low-redundancy sequence continuation — appears to train a *more direct, less self-doubting* reasoning style. Subsequent SFT reverses this: chains grow past base length (125.8 tokens), backtracking and hedging return. This is the same CPT-shapes / SFT-re-shapes signature seen on the knowledge axes, now visible in reasoning *process* rather than outcome — a effect invisible to accuracy alone. (it and it-bio again coincide, confirming the clean vocabulary ablation.)

**Table 3.** (4) Reasoning-behavior diagnostics on GSM8K (200 questions, *k*=5 samples, shared 8-shot CoT). CPT shortens and “decisifies” the chain without hurting accuracy; SFT reverses it.

| Checkpoint | Chain len (tok) | Backtrack/gen | Hedge/gen | Self-consist. | Acc. (2nd) |
| --- | --- | --- | --- | --- | --- |
| Base (it) | 108.8 | 0.370 | 0.001 | 0.749 | 0.667 |
| Base (it-bio) | 107.5 | 0.324 | 0.000 | 0.728 | 0.652 |
| **BioCPT** | **64.4** | **0.006** | 0.000 | 0.745 | 0.689 |
| BioCPT+SFT | 125.8 | 0.396 | 0.018 | 0.757 | 0.703 |

**Figure 2.**
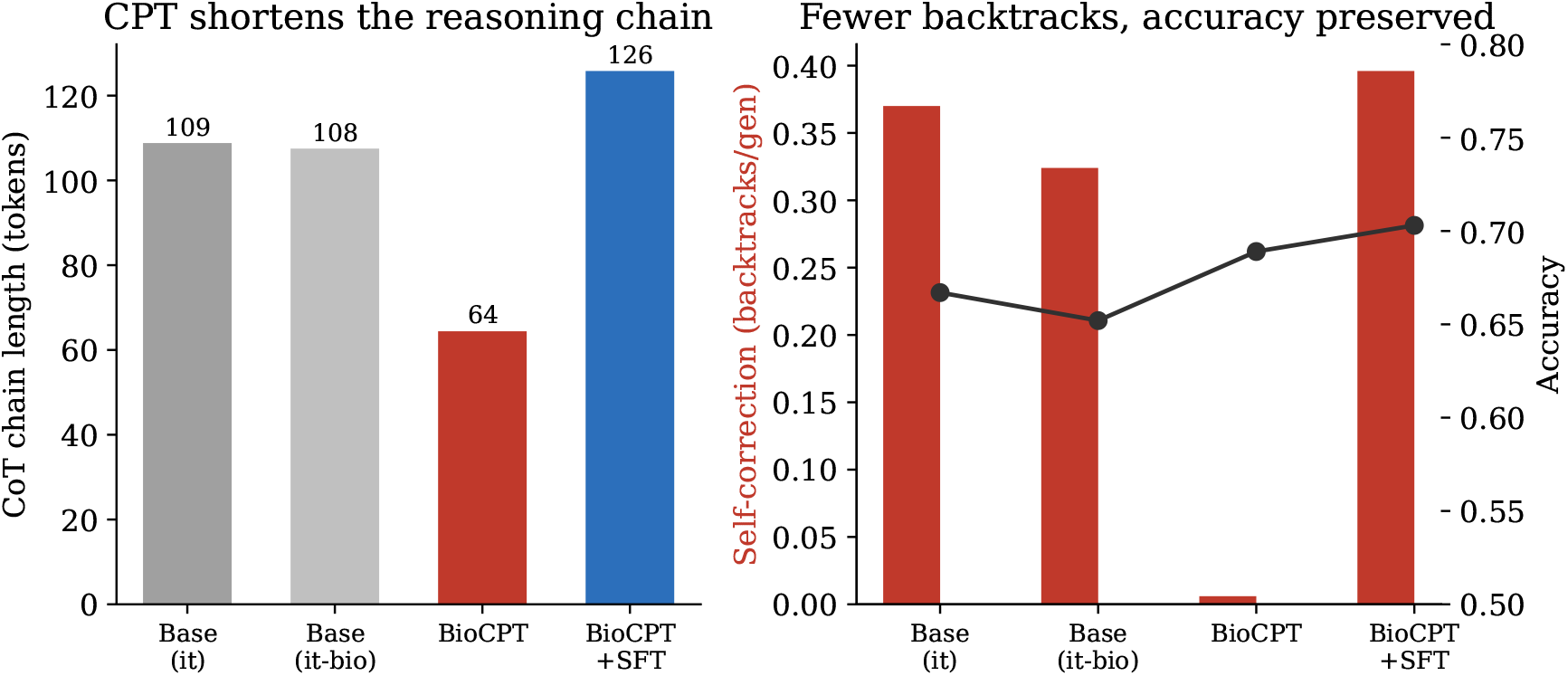
Reasoning behavior on GSM8K. **Left:** biological CPT shortens the chain-of-thought by 41%. **Right:** self-correction (backtracks) nearly vanishes at BioCPT while accuracy is preserved (even slightly improved); SFT reverses both. CPT instills a more concise, decisive reasoning style.

**Figure 3.**
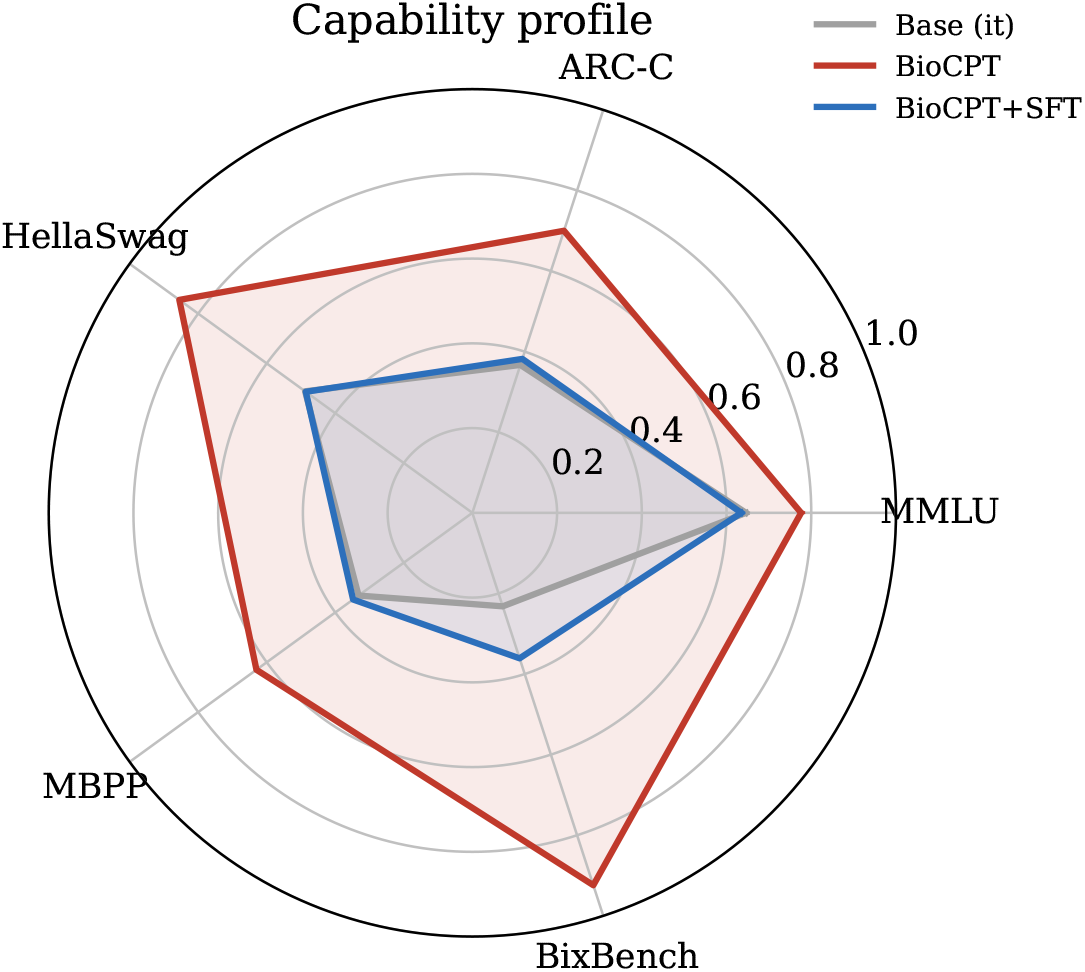
Capability profile of the three trained states. BioCPT (red) dominates the base model on every axis; SFT (blue) retreats toward base on all held-out axes. The biological CPT stage, not SFT, is what expands the profile.

### 3.6 Summary

Across every completed axis the ordering is identical: base *<* **BioCPT** *>* SFT, with BioCPT the peak (Figure 3). Biological CPT is not a tax; it is an investment in a shared capability substrate. SFT then trades breadth for task focus.

## 4 Discussion

### 4.1 A division of labor between CPT and SFT

The recurring shape of our results — base *<* BioCPT *>* SFT on every held-out axis — points to a division of labor that is usually conflated. Continued pretraining does not simply add domain facts; it *re-organizes and lifts a shared capability substrate* that is then visible on tasks (general knowledge, code) far from the pretraining domain. Supervised fine-tuning does the opposite: it *narrows* the model onto the target task distribution, trading breadth on held-out axes for focus on the served task. Neither stage is a substitute for the other, and budgeting them as interchangeable — or omitting CPT to “save compute” — misreads their roles. This is consistent with, and sharpens, prior router-level evidence on this model family that CPT reshapes middle-layer computation while SFT re-aligns the output.

### 4.2 Biological sequence as structured scientific data

If narrow, out-of-distribution biological sequence *raises* general and coding competence, then the “narrow domain = capacity competition” framing is inadequate. We propose instead that biological sequence functions as *structured scientific data*: a corpus whose value to a foundation model lies not (only) in the facts it carries but in the structural regularities it forces the model to internalize — regularities that transfer to other structured/formal languages such as code. Under this view biology is simply the first representative member of a broader class (code, mathematics, chemistry/SMILES), and the natural next question is not “does biology help biology” but “how does the *composition* of scientific-data CPT shape the capability profile of a foundation model.”

### 4.3 Limitations

This is a training-free re-analysis of one lineage of one 26B MoE model; the CPT / SFT recipe is fixed, so we characterize *this* lineage rather than establishing a scaling law. Part of the ARC/HellaSwag gain is a format effect (instruct-vs-completion loglikelihood), which we flag and which does not affect the knowledge-heavy MMLU gain. Zero-shot sequence-homology is not measurable by simple probes, so that axis is argued indirectly. TruthfulQA is a genuine, if small, regression and we do not hide it. Establishing the optimal CPT data mixture, and testing whether the CPT-lifts / SFT-narrows pattern generalizes across model sizes and domains, is the subject of the mixture-pretraining program we outline as future work.

## Data and code availability

All three (four) checkpoints derive from a public lineage of Gemma-4-26B-A4B; the merged BioCPT checkpoint is released as `dnagpt/OmniGene-4-CPT-v2-merged`. Evaluation scripts (general, coding, biology-loglikelihood, and chain-of-thought diagnostics), the exact few-shot configurations, and per-example model outputs for every table in this paper are released at the project repository. All benchmarks (MMLU, ARC, HellaSwag, TruthfulQA, MBPP, HumanEval, GSM8K, BixBench) are public; no proprietary data were used. The entire study is a training-free re-analysis and reproduces on a single 96 GB GPU.

## Acknowledgments Declaration of interests

The author declares no competing interests.

## Notes

https://github.com/maris205/gemma4-bio

### Competing Interest Statement

The authors have declared no competing interest.

